# Proteomic insight into soybean response to flooding stress reveals changes in basic energy metabolism and cell wall modifications

**DOI:** 10.1101/2021.04.08.438935

**Authors:** Mudassar Nawaz Khan, Iftikhar Ahmad, Israr-ud-Din, Essa Ali, Ahmed Noureldeen, Hadeer Darwish, Muhammad Ishfaq Khan, Muhammad Tayyab, Majid Khan

## Abstract

Soybean is a legume crop enriched with proteins and oil. It is frequently exposed to anthropogenic and natural flooding that limits its growth and yield. Current study applied gel-free proteomic techniques to unravel soybean response mechanism to flooding stress. Two-days-old soybeans were flooded for 4 days continuously and root samples were collected at days 2 to 6 for proteomic and enzymatic analyses. Age-matched untreated soybeans were collected as control. After protein extraction, purification and tryptic digestion, the peptides were analyzed on nano-liquid chromatography-mass spectrometry. A total of 539 and 472 proteins with matched peptides 2 or more were identified in control and flooded seedlings, respectively. Among these 364 proteins were commonly identified in both control and flooded soybeans. Fourty-two protein’s abundances were changed 4-fold after 2-days of flooding stress as compared to starting point. The cluster analysis showed that highly increased proteins included cupin family proteins, enolase, pectin methylesterase inhibitor, glyoxalase II, alcohol dehydrogenase and aldolase. The enzyme assay of enolase and pectin methylesterase inhibitor confirmed protein abundance changes. These findings suggest that soybean adopts the less energy consuming strategies and brings biochemical and structural changes in the cell wall to effectively respond to flooding stress and for the survival.

## Introduction

Soybean (*Glycine max* (L.) Merr.) is an important legume that is enriched with proteins and oil contents (Panizzi and Mandarino 1994). Frequent flooding due to climatic changes and ill-drained fields is one of the abiotic stresses that reduce its growth and yield (Githiri et al. 2006). Flooding initially causes damage to the roots (Sauter 2013), reduce the nutrient uptake (Sallam and Scott 1987) and decrease the nitrogen fixation capacity (Sung 1993). Flooding stress reduces biomass, tap-root length, and pod number, inhibits carbon/nitrogen content in root/nodule, decrease nodule dry weight, and grain yield in soybean (Miao et al. 2012). These reports suggest that flooding is a major constraint on growth and yield of soybean.

Root is an important primary organ to feel the effects of flooding stress. Flooding reduces the root dry weight first (Shimamura et al. 2003). Oxygen transport from the air to the roots is important for root physiology (Armstrong 1980). Flooding causes oxygen deficiency leading to hypoxia or anoxia as oxygen moves ten thousand times slower in water than in the air (Armstrong 1980; Armstrong and Drew, 2002). Plants respond to flooding stress by formation of adventitious roots (Shimamura et al. 2003; Mano and Omori 2007) and aerenchyma formation (Shimamura et al. 2003). Adventitious roots formation benefit the plant growth during flooding exposure (Rich et al. 2012). Flooding stress did not affect root growth of submergence-tolerant rice genotypes (Ismail et al. 2009). Roots undergo structural and functional alterations at the cellular, molecular and phenotypic level to deal with the flooding stress (Atkinson and Urwin 2012). Roots rapidly use starch reserves for limiting the damage and maintaining the growth (Sauter 2013).

Proteomic techniques found extensive applications in investigating effects of flooding stress and flooding stress-responsive proteins. Proteins belonging to the categories of glycolysis, fermentation, detoxification of reactive oxygen species, anaerobic catabolism, storage, stress, development, cell organization, transport, signaling and amino acid metabolism-related proteins were changed in abundance under flooding stress (Nanjo et al. 2010, 2013; Komatsu et al. 2012). Proteins related to the cell wall lignification were suppressed (Komatsu et al. 2010a). Protein abundances of energy-related proteins were raised whereas those involved in protein folding and cell structure organization were lowered in flooded soybean (Nanjo et al. 2012). Kamal et al. (2015) reported a decrease in sucrose metabolism-related proteins but increase in fermentation-related proteins in soybean cotyledon under flooding stress. Photosynthesis, RNA, DNA, signaling, and the tricarboxylic acid cycle were changed in abundance leaf, hypocotyl and root of soybean under flooding stress (Wang et al. 2017). Proteomics approaches have also been applied on subcellular level to reveal localized cellular responses and investigate communications among subcellular components during flooding stress. In the plasma membrane, proteins related to signaling, stress and the antioxidative system were increased; whereas, reactive-oxygen species scavenging enzymes activities were retarded in the cell wall (Komatsu et al. 2018). Protein metabolism-related proteins were decreased in the nucleus and also proteins related to electron transport chain were suppressed in the mitochondria (Komatsu et al. 2018). The soybean responses to flooding stress are being studied at various levels utilizing proteomic approaches. Current proteomic study was designed to analyze response mechanism of soybean to continuous four days flooding stress.

## Materials and Methods

### Plant material, growth conditions and treatment

Seeds of soybean (cv. Enrei) were sterilized with 2% sodium hypochlorite solution and washed in clean water. The sterilized seeds were sown 4 cm inside quartz sand in seedling cases (145 x 55 x 95 mm^3^) wetted with 150 mL water and grown at 25°C in a growth chamber (Sanyo, Tokyo, Japan) under fluorescent light (160 μmol m^-2^ s^-1^, 16 h light period/day). Eight seeds were grown in each pot per treatment. Two-day-old soybeans were flooded until day 6. The root samples were collected at days 2, 3, 4, 5 & 6 from un-treated control [labeled as 2(0), 3(0), 4(0), 5(0), 6(0)] and treated [labeled as 3(1), 4(2), 5(3), 6(4)] plants (Fig 1).

**Fig 1.**
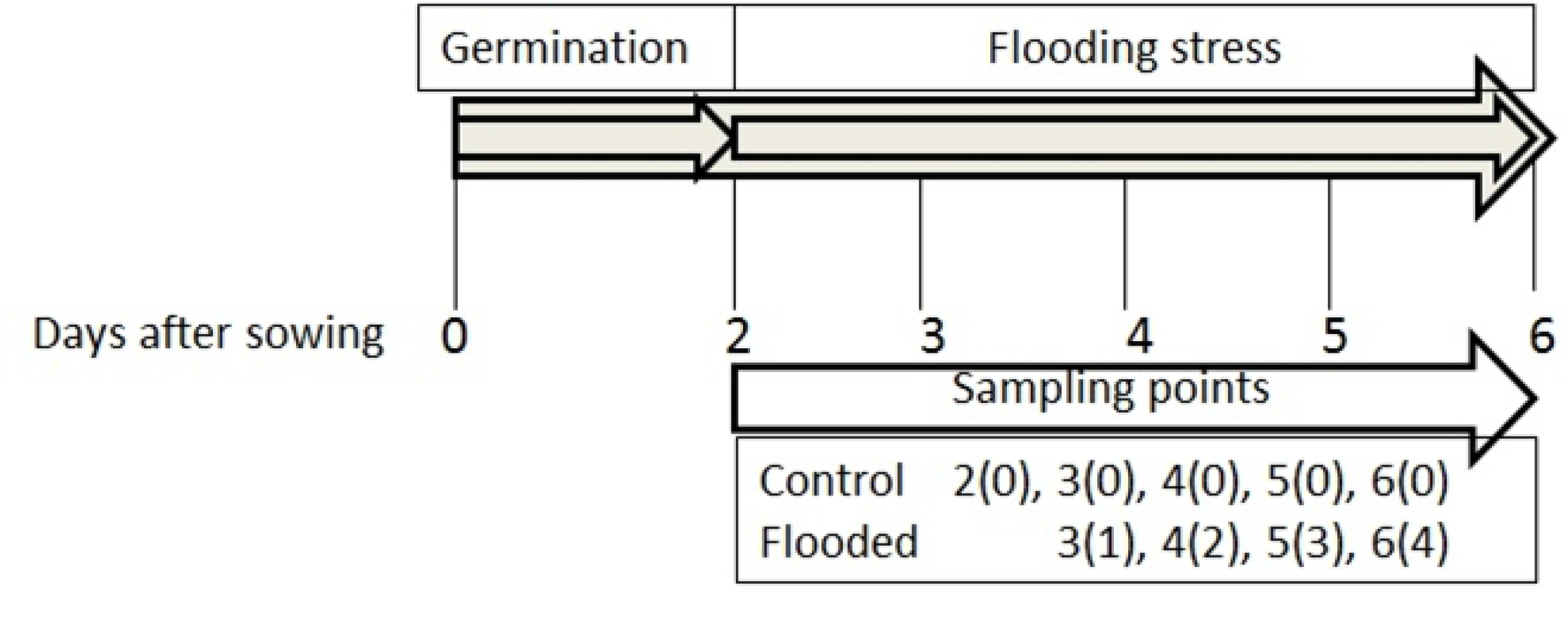
Experimental design of the study.

### Protein extraction

An amount of 500 mg of root was ground under liquid nitrogen using a mortar and pestle. The powder was transferred to an acetone solution containing 10% trichloroacetic acid and 0.07% 2-mercaptoethanol. The mixture was vortexed and sonicated for 10 min. The suspension was incubated for 1 h at -20°C and then centrifuged at 9,000×g at 4°C for 20 min. The pellet was washed twice with 0.07% 2-mercaptoethanol in acetone and dried. It was resuspended in lysis buffer (7 M urea, 2 M thiourea, 5% CHAPS, 2 mM tributylphosphine) by vortexing for 1 h at 25°C and centrifuged at 25°C with 20,000×g for 20 min. The supernatant was collected as protein extract. Bovine serum albumin was used as standard for protein concentration calculations through Bradford assay (Bradford et al. 1976).

### Protein purification and digestion for mass spectrometry analysis

Protein extracts of 100 μg were purified with methanol and chloroform to remove detergent from the samples. For purification and digestion of extracted proteins, methodology described by Khan and Komatsu (2016) was followed. The resulting tryptic peptides were acidified in 20% formate and analyzed by nano-liquid chromatography (LC) mass spectrometry (MS).

### Nanoliquid chromatography-tandem mass spectrometry analysis

A nanospray LTQ Orbitrap mass spectrometer (Thermo Fisher Scientific, San Jose, CA, USA) was operated in data-dependent acquisition mode with the installed XCalibur software (version 2.0.7, Thermo Fisher Scientific). The nanoLC-MS conditions and method as described by Khan and Komatsu (2016) was followed.

### Protein identification by Mascot search

Proteins were identified from a soybean peptide database constructed from the soybean genome database (Phytozome version 9.1, http://www.phytozome.net/soybean) (Schmutz et al. 2010) using the Mascot search engine (Matrix Science, London, UK). The data files were processed using Proteome Discoverer software (Thermo Fisher Scientific). The carbamidomethylation of cysteine was set as a fixed modification and oxidation of methionine was set as a variable modification. Trypsin was specified as the proteolytic enzyme and one missed cleavage was allowed. Peptide mass tolerance was set at 10 ppm and fragment mass tolerance was set at 0.8 Da.

### Differential analysis of acquired mass spectrometry data

The Mascot results were exported for SIEVE software analysis (version 2.1; Thermo Fisher Scientific). SIEVE compares the relative abundances of peptides and proteins between control and experimental groups. The MS detected peaks were aligned and the peptide peaks were detected as frames. Frames were generated for all parent ions scanned by MS/MS and were matched to exported Mascot results. In the differential analyses of protein profiles, total ion current was used as a normalization factor. For differential analyses, only proteins with at least two peptide matches across the data from all sample groups and replicates were defined as identified proteins.

### Cluster and *in silico* protein-protein interaction analyses

Protein ratios obtained from SIEVE software analysis were subjected to cluster analysis using Genesis software (version. 1.8.1; http://genome.tugraz.at) (Sturn et al. 2002). Cluster analysis was performed using hierarchical clustering with a Euclidean distance metric and a centroid linkage clustering method. The clustered proteins alignment in treatment was used for heat map generation in control. Clustered proteins were analyzed for *in silico* protein-protein interactions utilizing online STRING (version 11.0; https://string-db.org) program.

### Functional categorization

The functional categories of identified proteins were determined through MapMan bin codes using MapMan software (http://mapman.gabipd.org) (Usadel et al. 2005).

### Analysis of enzyme activities Enolase

A quantity of 200 mg of root was homogenized in lysis buffer (20 mM Tris-HCl pH 7.5, 1 mM EDTA, 1 mM 2-mercaptoethanol). The suspension was centrifugation at 20,000×g at 4°C for 30 min. Protein concentrations were estimated by Bradford assay (Bradford 1976). A reaction mixture consisting of 100 mM triethanolamine (pH 7.4), 120 mM KCl, 2.25 mM 2-phosphoglycerate, 0.2 mM 2-NADH, 30 mM MgSO_4_, 1.75 mM ADP, 10 units pyruvate kinase, and 15 units L-lactic dehydrogenase was used for enzymatic assay. Enzyme extract of 100 μL was mixed with 900 μL of reaction mixture and vortexed. The absorbance was measured at 340 nm using a UV/Vis spectrophotometer (Anderson et al. 1984; Joseph et al. 1996).

### Plant invertase/pectin methylesterase inhibitor superfamily

Plant invertase assay was performed by slightly modifying protocol of Huang et al. (1998). The extraction procedure was performed on ice. A weight of 200 mg of soybean roots was used for enzyme extraction. Roots were ground into fine powder in liquid nitrogen and extracted in buffer that consisted of 50 mM HEPES-KOH, pH 7.4, containing 5% Polyvinyl pyrrolidone, 1 mM EDTA, 1 mM EGTA, 1 mM PMSF, 5 mM DTT, 0.1% Triton X-100, and 1% glycerol. The homogenate was centrifuged for 20 min at 15000×*g* in a refrigerated centrifuge. The supernatant was collected as the enzyme crude extract. The crude extract was vacuum-filtered through bottle-top vacuum filters (pore size: 0.45 μm). The filtrate was concentrated to about one-third of the volume by centrifuging for 45 min at 2000×*g*. The supernatant was used for enzyme assay. An enzyme extract of 100 μL was mixed with 900 μL of reaction mixture and reduction in absorbance was measured at 340 nm using a UV/vis spectrophotometer.

### Statistical analysis

Enolase and Pectin methylesterase activities were analyzed for statistical significance using Duncan’s multiple comparison test p<0.05.

## Results

### Identified proteins in soybean root under flooding stress

To identify differentially changed proteins in soybean root, a gel-free proteomic technique was used to analyze the protein profiles of soybeans that had been flooded continuously for 4 days. A total of 539 and 472 proteins with matched peptides 2 or more were identified in control (S1 Table) and flooding-stressed soybean roots (S2 Table), respectively. Out of the total identified proteins, 364 were commonly identified in control and flooding-stressed plants (S3 Table; Fig 2). Among these 364 proteins, protein abundances of 42 proteins were changed 4-fold in flooding-stressed plants after 2-days of flooding (Table 1).

**Fig 2.**
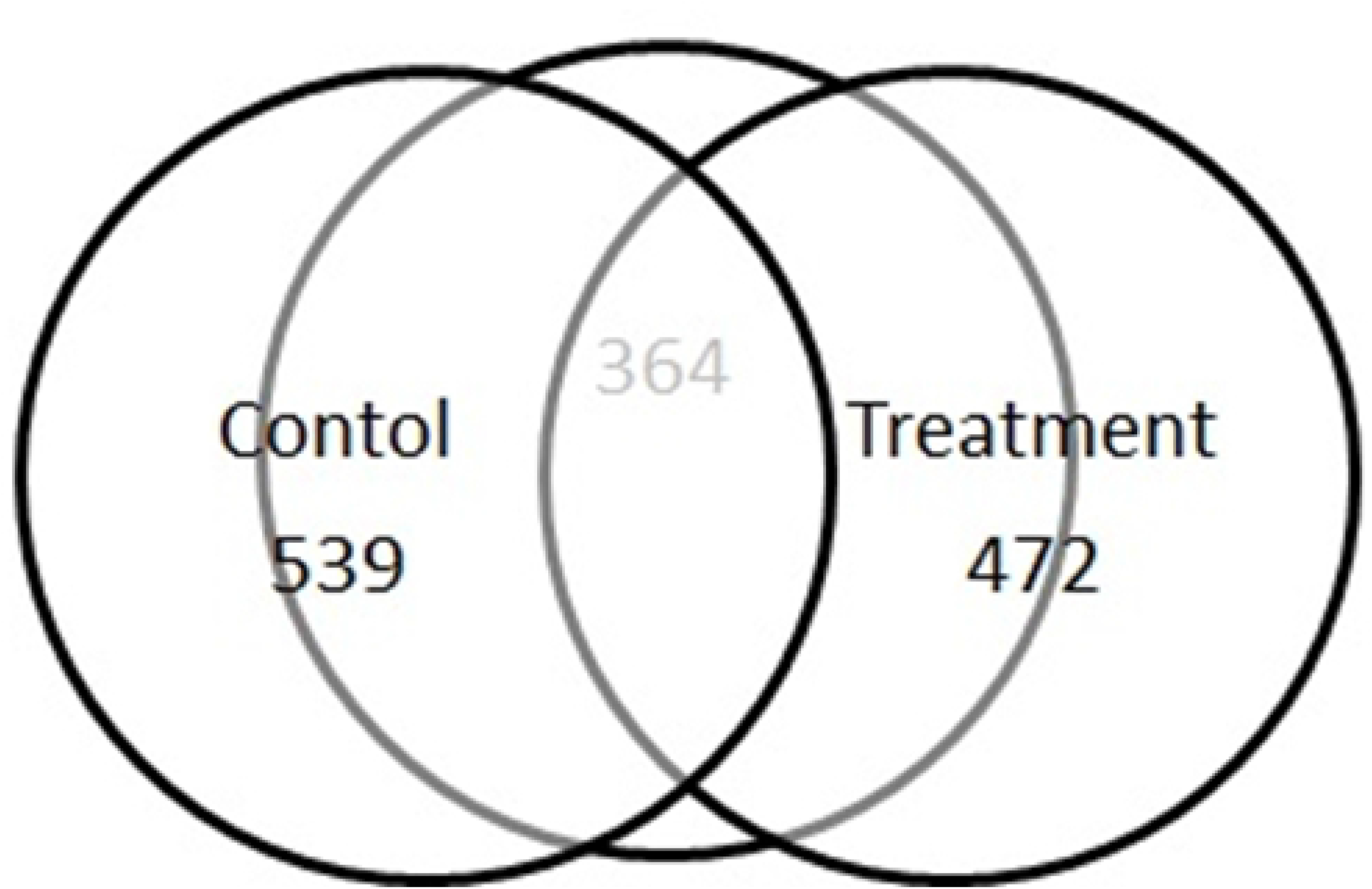
Venn diagram of total identified and common proteins in control and flooding-stressed soybean seedlings.

**Table 1.** Proteins identified in soybean that changed 4-folds in abundance after 2 days flooding as compared to starting point 2(0) *.

| Protein ID | Description | Peptides | Protein abundance Ratios for Control soybean |  |  |  | Protein abundance Ratios for flooding-stressed soybean |  |  |  | Functional Category |
| --- | --- | --- | --- | --- | --- | --- | --- | --- | --- | --- | --- |
|  |  |  | 3(0)/2(0) | 4(0)/2(0) | 5(0)/2(0) | 6(0)/2(0) | 3(1)/2(0) | 4(2)/2(0) | 5(3)/2(0) | 6(4)/2(0) |  |
| Glyma20g28466.1 | Cupin family protein | 2 | 0.55 | 0.96 | 0.22 | 0.41 | 0.19 | 57.06 | 69.19 | 0.60 | Development |
| Glyma03g03460.1 | Plant invertase/pectin methylesterase inhibitor superfamily protein | 2 | 5.15 | 26.74 | 9.96 | 38.39 | 1.57 | 44.03 | 49.65 | 28.87 | Cell wall |
| Glyma20g28550.1 | Seed maturation protein | 2 | 0.15 | 0.27 | 0.12 | 0.12 | 0.16 | 19.36 | 8.15 | 0.30 | Development |
| Glyma10g33350.2 | Arabidopsis thaliana peroxxygenase 2 | 3 | 0.72 | 0.20 | 0.26 | 0.17 | 0.29 | 12.04 | 1.79 | 0.54 | Development |
| Glyma03g07470.1 | Stress induced protein | 3 | 0.95 | 0.20 | 0.15 | 0.13 | 0.25 | 11.24 | 0.09 | 0.85 | Hormone metabolism |
| Glyma10g03310.1 | Seed maturation protein | 5 | 0.27 | 0.65 | 0.03 | 0.42 | 0.16 | 11.15 | 4.16 | 0.28 | Development |
| Glyma16g32960.1 | Enolase | 2 | 0.93 | 3.71 | 0.59 | 1.40 | 1.05 | 10.28 | 5.12 | 2.84 | Glycolysis |
| Glyma08g23750.4 | Ribosomal protein L30/L7 family protein | 4 | 2.89 | 12.42 | 1.08 | 3.29 | 0.07 | 9.61 | 3.20 | 0.14 | Protein |
| Glyma19g34780.1 | RmlC_like cupins superfamily protein | 7 | 0.63 | 0.80 | 0.24 | 0.56 | 0.07 | 9.19 | 2.98 | 0.51 | Development |
| Glyma11g15870.1 | RmlC_like cupins superfamily protein | 7 | 0.24 | 0.03 | 0.07 | 0.01 | 0.19 | 9.07 | 3.54 | 0.01 | Development |
| Glyma13g21291.1 | embryonic cell protein 63 | 6 | 0.89 | 0.49 | 0.29 | 0.13 | 0.18 | 9.07 | 0.30 | 0.72 | Development |
| Glyma11g02410.1 | RNA binding Plectin/S10 domain_containing protein | 2 | 1.04 | 4.19 | 2.20 | 2.57 | 0.14 | 8.05 | 2.78 | 0.42 | Protein |
| Glyma13g18450.2 | RmlC_like cupins superfamily protein | 9 | 0.22 | 0.00 | 0.06 | 0.05 | 0.02 | 7.92 | 0.13 | 0.02 | Development |
| Glyma20g28640.1 | Cupin family protein | 18 | 0.47 | 1.49 | 0.11 | 0.31 | 0.06 | 7.84 | 2.70 | 0.83 | Development |
| Glyma13g33590.1 | Glyoxalase II 3 | 5 | 1.32 | 2.01 | 1.35 | 2.16 | 0.40 | 7.40 | 6.06 | 2.60 | Biodegradation of Xenobiotics |
| Glyma13g17980.1 | Late embryogenesis abundant domain_containing protein / LEA domain_containing protein | 4 | 0.73 | 0.31 | 0.18 | 0.06 | 0.11 | 7.07 | 0.24 | 1.04 | Not assigned |
| Glyma12g06950.1 | Pathogenesis_related thaumatin superfamily protein | 2 | 0.83 | 0.72 | 0.37 | 0.85 | 0.31 | 6.87 | 3.21 | 0.12 | Stress |
| Glyma08g15000.1 | Ribosomal protein L6 family protein | 5 | 1.17 | 2.98 | 0.19 | 0.75 | 0.13 | 6.59 | 0.97 | 0.29 | Protein |
| Glyma09g02790.1 | Ribosomal protein L13 family protein | 2 | 3.42 | 11.87 | 0.61 | 2.36 | 0.12 | 5.88 | 2.30 | 0.91 | Protein |
| Glyma13g44261.1 | Cystathionine beta_synthase (CBS) protein | 3 | 0.41 | 0.23 | 0.05 | 0.16 | 0.18 | 5.85 | 2.49 | 0.62 | Not assigned |
| Glyma06g11940.1 | Ribosomal protein S3Ae | 4 | 0.75 | 5.78 | 0.51 | 0.64 | 0.12 | 5.84 | 0.26 | 0.08 | Protein |
| Glyma14g36620.1 | Ribosomal protein L16p/L10e family protein | 2 | 0.96 | 8.03 | 1.90 | 2.97 | 0.32 | 5.81 | 1.61 | 0.49 | Protein |
| Glyma12g11130.1 | beta_amylase 5 | 7 | 0.58 | 0.43 | 0.22 | 0.45 | 0.01 | 5.70 | 2.59 | 0.37 | Major CHO metab. |
| Glyma20g21370.1 | Ribosomal protein S13A | 2 | 1.80 | 3.94 | 0.72 | 1.64 | 0.26 | 5.50 | 1.30 | 0.38 | Protein |
| Glyma10g36880.4 | Ribosomal protein S13/S18 family | 3 | 1.11 | 3.18 | 0.78 | 1.94 | 0.02 | 5.17 | 0.96 | 0.10 | Protein |
| Glyma09g16606.1 | Ribosomal L22e protein family | 2 | 1.32 | 4.56 | 0.53 | 1.83 | 0.10 | 4.90 | 1.41 | 0.28 | Protein |
| Glyma16g23730.1 | Ribosomal protein S4 (RPS4A) family protein | 5 | 0.80 | 6.24 | 0.49 | 1.44 | 0.08 | 4.87 | 1.22 | 0.04 | Protein |
| Glyma10g39150.1 | Cupin family protein | 10 | 0.44 | 0.35 | 0.29 | 0.28 | 0.51 | 4.79 | 0.22 | 0.37 | Development |
| Glyma17g13760.1 | Adenylate kinase 1 | 3 | 0.91 | 3.37 | 1.45 | 2.13 | 0.02 | 4.74 | 1.84 | 1.24 | Nucleotide metab. |
| Glyma06g12780.1 | Alcohol dehydrogenase I | 6 | 0.67 | 1.26 | 0.54 | 0.96 | 0.28 | 4.68 | 3.77 | 1.51 | Fermentation |
| Glyma15g20180.1 | Sucrose synthase 4 | 6 | 0.45 | 2.68 | 0.98 | 2.00 | 0.20 | 4.68 | 1.61 | 0.37 | Major CHO metab. |
| Glyma14g34740.1 | Annexin 2 | 3 | 0.56 | 0.39 | 0.12 | 0.76 | 0.02 | 4.65 | 0.33 | 0.86 | Cell |
| Glyma03g32020.3 | RmlC_like cupins superfamily protein | 8 | 0.56 | 0.99 | 0.04 | 0.35 | 0.18 | 4.65 | 8.27 | 0.02 | Development |
| Glyma09g16553.1 | Ribosomal L22e protein family | 2 | 1.66 | 3.97 | 0.94 | 2.11 | 0.27 | 4.46 | 1.76 | 1.43 | Protein |
| Glyma11g00890.1 | Ribosomal protein S3Ae | 3 | 0.73 | 5.56 | 0.42 | 0.37 | 0.14 | 4.38 | 0.92 | 0.09 | Protein |
| Glyma08g08970.1 | Urease accessory protein G | 3 | 0.50 | 0.73 | 0.19 | 0.28 | 0.21 | 4.37 | 1.04 | 0.06 | Amino acid metab. |
| Glyma20g17440.1 | Uricase / urate oxidase / nodulin 35 putative | 3 | 0.50 | 1.26 | 0.24 | 1.22 | 0.65 | 4.33 | 3.85 | 0.23 | Nucleotide metab. |
| Glyma02g38730.1 | Aldolase superfamily protein | 3 | 0.77 | 2.30 | 0.57 | 1.32 | 0.63 | 4.25 | 3.22 | 0.55 | Glycolysis |
| Glyma17g22161.1 | Ribosomal protein S4 (RPS4A) family protein | 2 | 1.50 | 6.64 | 0.64 | 1.71 | 0.21 | 4.17 | 1.38 | 0.19 | Protein |
| Glyma17g10710.1 | Ribosomal protein S4 | 4 | 1.20 | 3.23 | 0.84 | 1.63 | 0.14 | 4.11 | 0.86 | 0.23 | Protein |
| Glyma19g01210.1 | Formate dehydrogenase | 2 | 0.72 | 0.85 | 0.13 | 1.06 | 0.52 | 4.07 | 2.47 | 0.08 | C1-metabolism |
| Glyma17g34070.1 | Class II aminoacyl_tRNA and biotin synthetases superfamily protein | 4 | 0.57 | 3.27 | 0.29 | 0.67 | 0.02 | 4.06 | 0.94 | 0.04 | Protein |

\*Starting point 2(0) is 1 and is used for abundance ratios calculation in both control and flooded seedlings.

### Identified proteins belonged to diverse functional categories

The total identified proteins in control (539) and flooded soybean (472) had 364 commonly changed proteins. The total identified proteins were functionally categorized according to MapMan codes (Fig 3). Maximum number belonged to ‘protein’ category with 152 in control and 117 in flooded soybeans. Proteins belonging to protein-metabolism-related category in-turn belonged to protein synthesis, degradation, folding and other related functions. The second major category was stress-related proteins with 33 identified in control and 34 in flooded seedlings. The other differentially changed proteins belonged to glycolysis (31 in control, 24 in flooded), amino acid metabolism (27 in both control & flooded), cell (25 in control, 22 in flooded), TCA/organic transformation (20 in control, 12 in flooded), signaling (20 in control, 18 in flooded), secondary metabolism (18 in control, 14 in flooded), development (18 in control, 23 in flooded), redox (17 in control, 19 in flooded), cell wall (17 in control, 16 in flooded), hormone metabolism (16 both in control & flooded), RNA (14 in control, 9 in flooded), transport (12 in control, 11 in flooded), mitochondrial electron transport (10 in control, 05 in flooded), lipid metabolism (9 in control, 5 in flooded), major CHO metabolism (8 in control, 10 in flooded), mitochondrial metabolism (7 in control, 6 in flooded) and fermentation (7 in control, 8 in flooded). The 25 proteins in control and 16 in flooded belonged to miscellaneous; while 22 in control and 31 proteins in flooded seedlings were not assigned any function. The ‘Others’ category included proteins related to organo-pentose phosphate pathway, C1-metabolism, minor carbohydrate metabolism, DNA, metal handling, biodegradation of xenobiotics, cofactor and vitamin metabolism, and photosynthesis.

**Fig 3.**
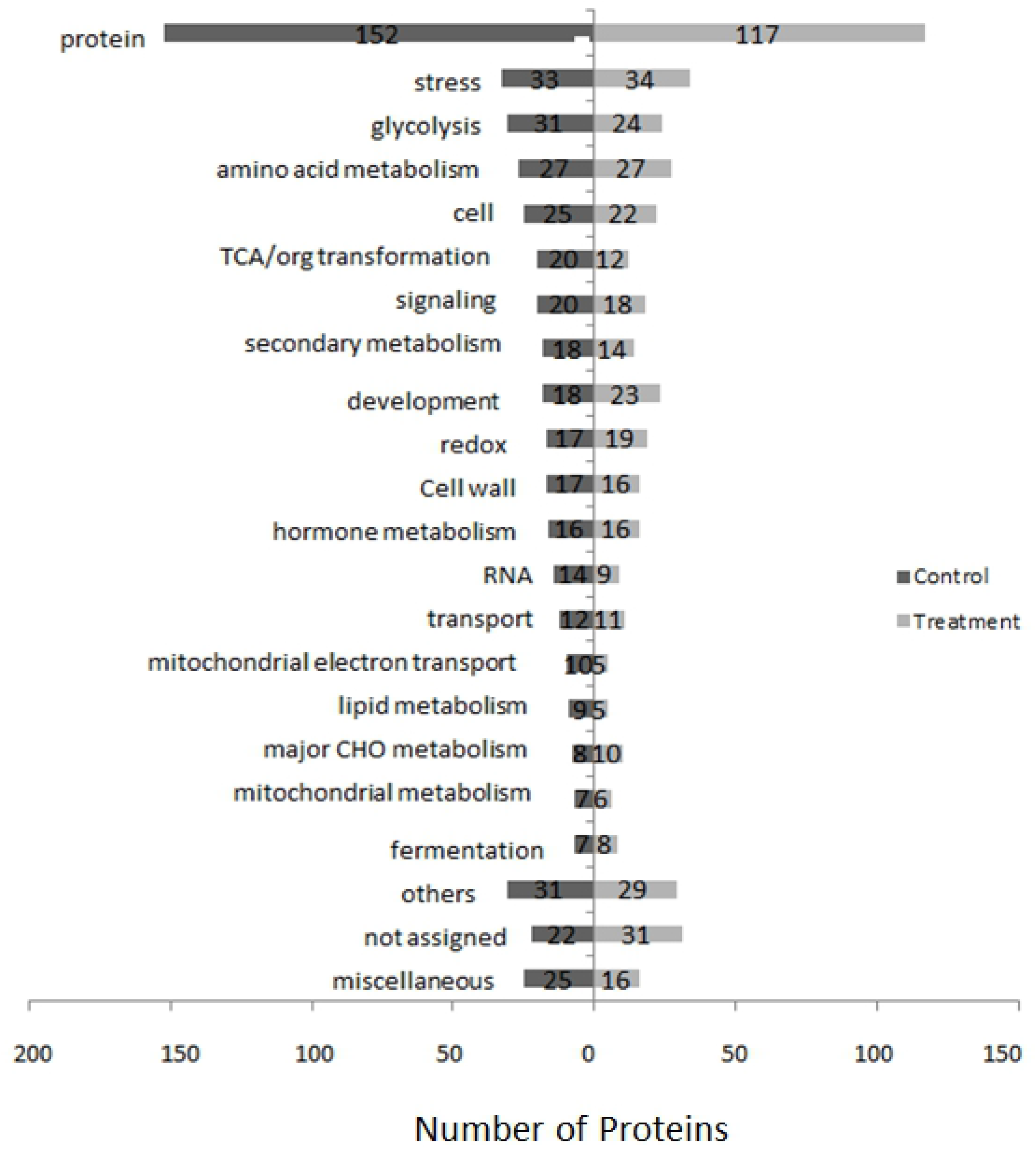
MapMan-based functional categorization of proteins identified in soybean roots exposed to flooding stress.

### High changes in protein abundances observed in soybean root under flooding stress

Among the total identified proteins in flooded and control soybeans, 42 common proteins increased in abundance 4-fold or more after 2-days flooding stress as compared to 2-days-old seedlings. The protein abundance changes in flooded plant proteins ranged from 4.06 to 57.06 fold when analyzed at 4(2). These proteins were subjected to cluster analysis that grouped protein abundance changes in flooded plants into 3 clusters (Fig 4A). In the first cluster, protein abundance of majority of proteins was increased at 2^nd^, 3^rd^ and 4^th^ day of flooding. Abundances of few proteins fell to the starting point at the end of 4-days flooding while a very few decreased. Cluster I contained 16 proteins that included cupin family protein (Glyma20g28466.1 & Glyma20g28640.1), plant invertase/pectin methylesterase inhibitor superfamily protein (Glyma03g03460.1), *Arabidopsis thaliana* peroxygenase 2 (Glyma10g33350.2), seed maturation protein (Glyma20g28550.1 & Glyma10g03310.1), RNA binding Plectin/S10 domain containing protein (Glyma11g02410.1), glyoxalase II 3 (Glyma13g33590.1), ribosomal protein L13 family protein (Glyma09g02790.1), ribosomal L22e protein family (Glyma09g16553.1), enolase (Glyma16g32960.1), RmlC like cupins superfamily protein (Glyma19g34780.1), cystathionine beta synthase (CBS) protein (Glyma13g44261.1), ribosomal protein L16p/L10e family protein (Glyma14g36620.1), alcohol dehydrogenase 1 (Glyma06g12780.1) and aldolase superfamily protein (Glyma02g38730.1).

**Fig 4.**
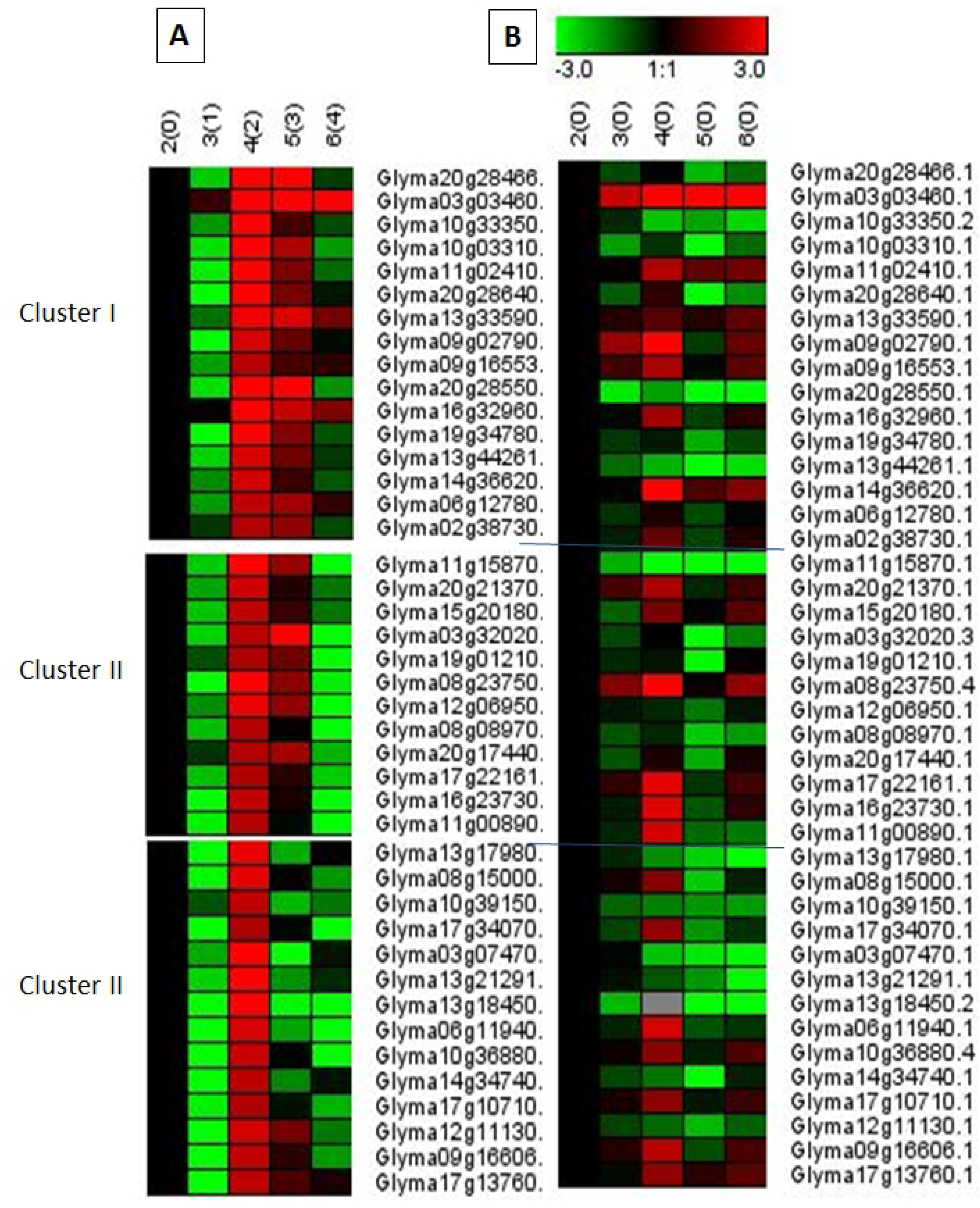
Cluster analysis of flooding-responsive proteins in flooded (A) and control (B) soybean roots using Genesis software.

In cluster II, protein abundance was increased until 3^rd^ day of flooding 5(3), but decreased even than the starting point 2(0) on the next day. The proteins grouped in the 2^nd^ cluster included RmlC like cupins superfamily protein (Glyma03g32020.3 & Glyma11g15870.1), ribosomal protein S13A (Glyma20g21370.1), sucrose synthase 4 (Glyma15g20180.1), formate dehydrogenase (Glyma19g01210.1), ribosomal protein L30/L7 family protein (Glyma08g23750.4), Pathogenesis-related thaumatin superfamily protein (Glyma12g06950.1), urease accessory protein G (Glyma08g08970.1), uricase/urate oxidase/nodulin 35 putative (Glyma20g17440.1), ribosomal protein S4 (RPS4A) family protein (Glyma16g23730.1 & Glyma17g22161.1) and ribosomal protein S3Ae (Glyma11g00890.1).

In cluster III, protein abundance was increased four-fold at 3^rd^ day of flooding 5(3), but decreased for majority of proteins in the next 2 days of flooding. The proteins grouped in the 3^rd^ cluster included late embryogenesis abundant domain containing protein/LEA domain containing protein (Glyma13g17980.1), ribosomal protein L6 family protein (Glyma08g15000.1), cupin family protein (Glyma10g39150.1), Class II aminoacyl tRNA and biotin synthetases superfamily protein (Glyma17g34070.1), stress induced protein (Glyma03g07470.1), embryonic cell protein 63 (Glyma13g21291.1), RmlC like cupins superfamily protein (Glyma13g18450.2), ribosomal protein S3Ae (Glyma06g11940.1), ribosomal protein S13/S18 family (Glyma10g36880.4), annexin 2 (Glyma14g34740.1), ribosomal protein S4 (Glyma17g10710.1), beta amylase 5 (Glyma12g11130.1), ribosomal L22e protein family (Glyma09g16606.1) and adenylate kinase 1 (Glyma17g13760.1).

In control plants, these proteins were aligned to check abundance changes (Fig 4B). Control plant proteins aligned against flooded cluster I revealed different pattern of abundance changes except for the plant invertase. The protein abundances of *Arabidopsis thaliana* peroxygenase 2, seed maturation protein, cupin family protein, glyoxalase II 3, enolase, RmlC like cupins superfamily protein, cystathionine beta synthase protein, alcohol dehydrogenase 1 and aldolase superfamily protein were decreased in control as compared to same-aged flooded plants. In control plant proteins aligned against flooded cluster II, abundances of RmlC like cupins superfamily protein, formate dehydrogenase, and urease accessory protein G were very less as compared to age-matched flooded plants. In control plant proteins aligned against flooded cluster III, LEA domain containing protein, cupin family protein, stress induced protein, embryonic cell protein 63, RmlC like cupins superfamily protein, annexin 2 and beta amylase 5 were decreased in abundance throughout the growth period; whereas, these proteins were increased in flooded plants.

### Compact Protein-protein interactions revealed under flooding stress

*In silico* Protein-protein interactions were estimated by using STRING (version 11.0) (Fig 5). Among the 42 common proteins, 14 proteins were found to strongly interact with each other forming a complex network. These included ribosomal protein S4 family protein (Glyma16g23730.1), ribosomal protein L16p/L10e family protein (Glyma14g36620.1), ribosomal protein S3Ae (Glyma11g00890.1, Glyma06g11940.1), ribosomal protein S13A (Glyma20g21370.1), ribosomal protein S4 (Glyma17g10710.1, Glyma17g22161.1), ribosomal L22e protein family (Glyma09g16553.1, Glyma09g16606.1), ribosomal protein L6 family protein (Glyma08g15000.1), ribosomal protein S13/S18 family (Glyma10g36880.4), ribosomal protein L30/L7 family protein (Glyma08g23750.4), ribosomal protein L13 family protein (Glyma09g02790.1), and RNA binding Plectin/S10 domain containing protein (Glyma11g02410.1). Lesser interacting proteins included cupin family protein (Glyma10g39150.1, Glyma20g28640.1), RmlC like cupins superfamily protein (Glyma13g18450.2, Glyma11g15870.1), embryonic cell protein 63 (Glyma13g21291.1), and seed maturation protein (Glyma20g28550.1). Some other proteins were not found to interact with each other as can be seen isolated in the figure 4.

**Fig 5.**
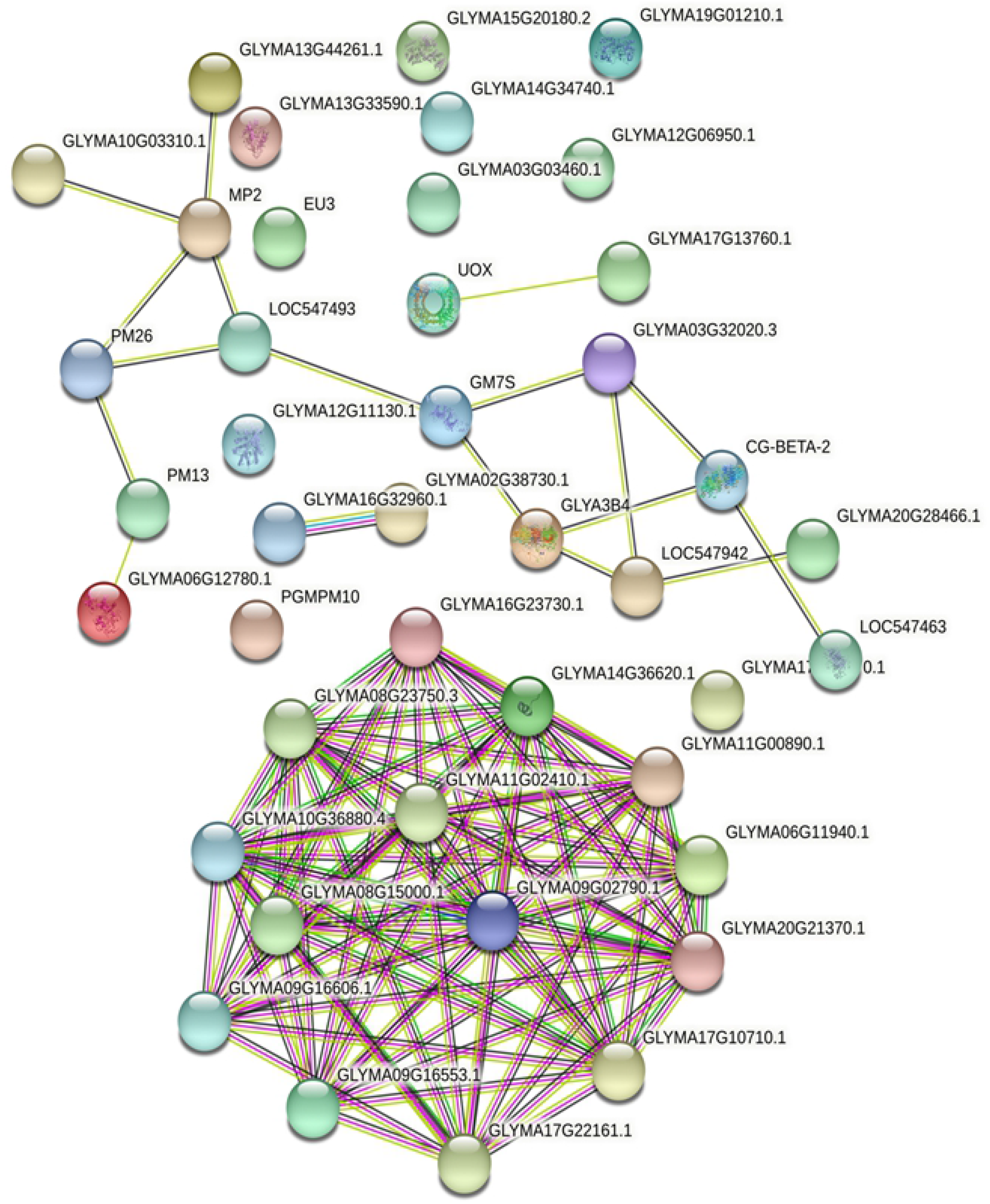
Protein-protein interactions network among the differentially changed proteins analyzed through STRING.

### Enolase and Plant invertase/pectin methylesterase inhibitor show highly significant response to flooding stress

The enzyme enolase which is also called phosphopyruvate hydratase is an important enzyme of glycolysis was analyzed for activity changes under flooding stress. The protein abundance of enolase was highly increased under initial 2 days of flooding stress (10.28) and decreased gradually latter at day 3 and 4 of flooding stress (5.82 & 2.84) (Fig 6A). While in control plants, there was no appreciable increase with increasing age. The results of enolase activity assay followed the pattern of protein abundance. The enzyme activity tremendously increased from first to second day of flooding (160.65 to 720.15 unit/mg protein) and gradually decreased at day 3 and 4 of flooding (600.25 & 470.58 unit/mg protein, respectively) (Fig 6B). The changes in activity were significant as compared to those observed in control plants and also among the different flooding duration.

**Fig 6.**
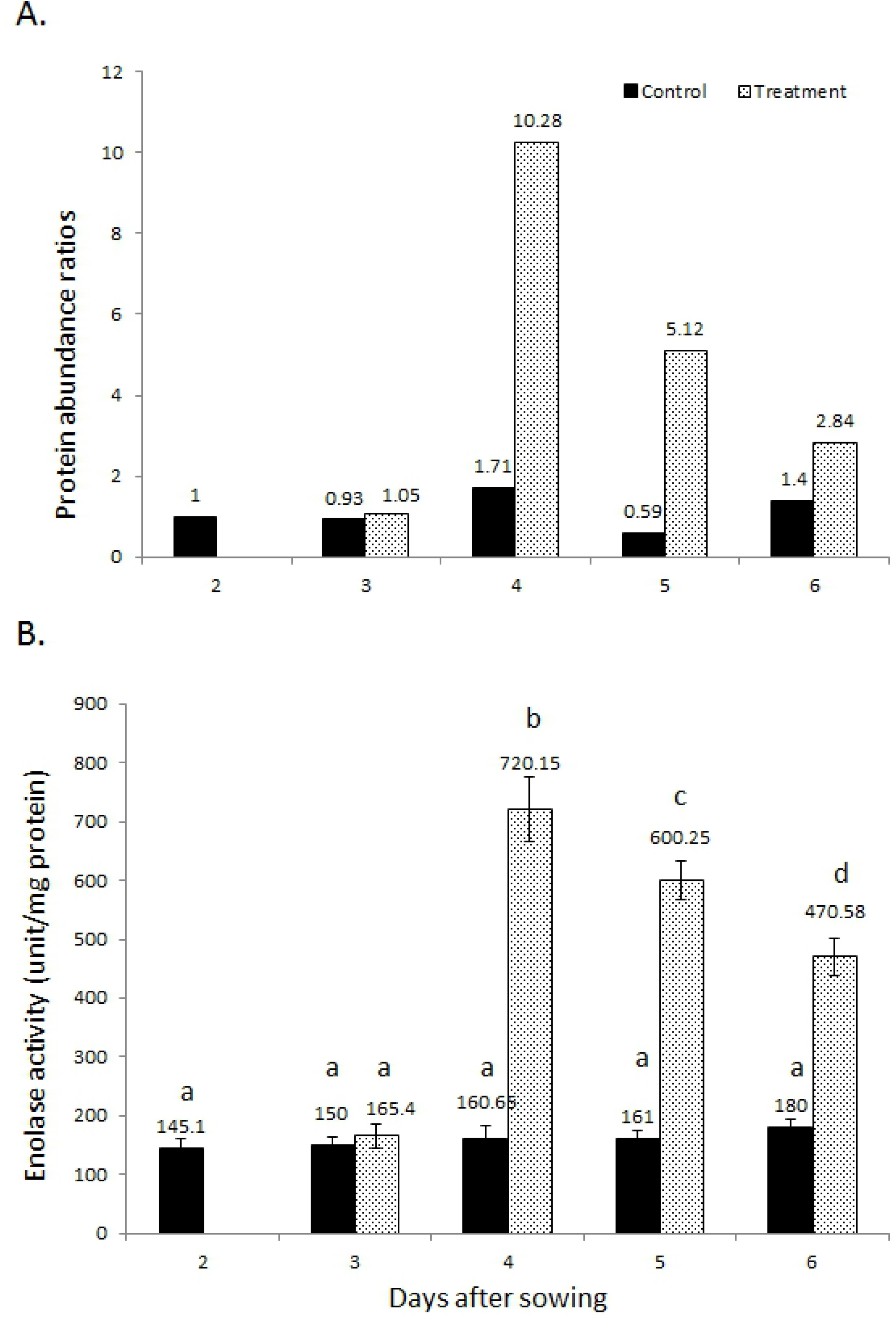
Changes in protein abundance (A) and enzyme activity (B) of enolase in soybean roots under flooding stress. Different alphabets indicate significant changes.

Plant invertase also called pectin methylesterase inhibitor (PMEI) showed a high increase in protein abundance (Fig 7A). The protein abundance increased from 1.57 after 1 day of flooding towards maximum of 49.65 at the end of 3 days flooding. It deceased at the end of 4 day of flooding to a level of 28.87. The enzyme activity of plant invertase was analyzed in control and flooded plants (Fig 7B). PMEI activity gradually increased 90.77 at 1 day flooding to a highest of 390.47 unit/mg protein at the end of 4-days flooding period. The activity changes were statistically significant in the last 2 days of flooding.

**Fig 7.**
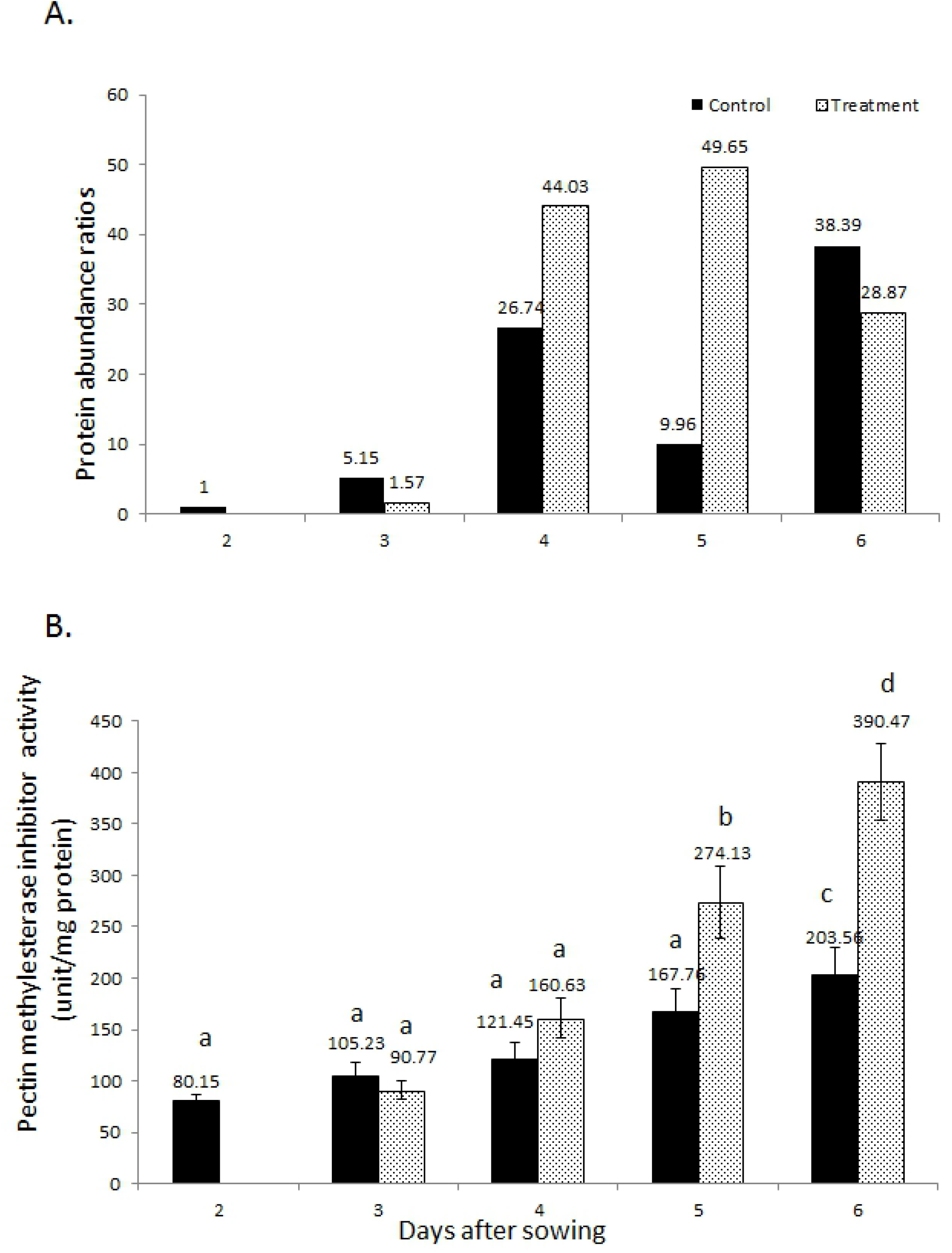
Changes in protein abundance (A) and enzyme activity (B) of plant invertase/pectin methylesterase inhibitor in soybean roots under flooding stress. Different alphabets indicate significant changes.

## Discussion

Flooding stress causes injury in the soybean (Komatsu et al. 2012). In the current study, continuous flooding stress was applied to the soybeans for 4 days and protein abundance changes were analyzed through gel-free proteomic technique. The study was conducted to unravel the mechanism involved in soybean responses to continuous flooding stress. Flooding stress brought huge abundance changes in many physiologically important proteins. Among the functionally important proteins, abundances of cupin family protein, RmlC like cupins superfamily protein, enolase, plant invertase/pectin methylesterase inhibito protein, *Arabidopsis thaliana* peroxygenase 2, seed maturation protein, glyoxalase II 3, alcohol dehydrogenase 1 and aldolase supefamily protein were significantly increased under flooding stress as compared to starting point 2(0) as well as control plants.

RmlC-like cupin superfamily proteins and cupin family proteins, which include storage proteins belonging to the development category, were highly increased in abundance under flooding stress. Cupin are functionally very diverse family of proteins (Dunwell et al. 2004) and play role in seedling development in soybean (Lapik and Kaufman, 2003). Cupins and seed maturation proteins with nutrient reservoir activity, are development-related storage proteins that were also previously reported to be increased in flooded soybean roots possibly due to delayed degradation (Salavati et al. 2012; Komatsu et al. 2010b). The results of the current study suggest delayed use of cupins as storage proteins in the initial 3 days of flooding stress as against control plants where their abundance was quite low. The other types of cupins modify the structure of cell wall as phosphomannose isomerase modifies mannose derivatives (Nunez et al. 2000). Cupins such as dTDP-rhamnose enzymes produce activated rhamnose as germin cross-link the plant cell-wall components (Giraud et al. 2000; Ma et al. 2001). Hence cupins are vital for cell survival through modification of cell wall. The increased abundance of cupins in the flooded soybean may point out towards their role in maintaining cell wall integrity under flooding stress.

Glyoxalase II was increased in flooded 7-fold as compared to starting point and 3-fold as compared to 4-days age-matched control. This enzyme is involved in detoxification of methylglyoxal whose production is increased many-folds under abiotic stress (Yadav et al. 2005). Methylglyoxal II is produced as by-product of metabolic pathways such as glycolysis and from photosynthesis intermediates (glyceraldehyde-3-phosphate & dihydroxyacetone phosphate). Methylglyoxal is a reactive cytotoxin that can cause lipid peroxidation, oxidation of proteins & fatty acids and disruption of membranes (Chaplen, 1998; Gill and Tuteja, 2010). Methylglyoxal is detoxified by glyoxalase system consisting of glyoxalase I and glyoxalase II that catalyze conversion of methylglyoxal to D-lactate while using glutathione as co-factor (Yadav et al. 2005). The increased protein abundance of glyoxalase II in current study showed an increase in detoxification of methylglyoxal as a defense effort by soybean.

Aldolase superfamily protein abundance was increased at 2^nd^ and 3^rd^ days of flooding as compared to control plants. Aldolase enzyme is an enzyme that brings conversion of fructose bisphosphate to glyceraldehyde-3-phosphate and dihydroxyacetone phospate, an important step of glycolysis. The enzyme is also involved in gluconeogenesis and calvin cycle (e (Rutter, 1964; Berg et al. 2010). Nuclear isoform of fructose-bisphosphate aldolase regulates expression of its own gene as well as other genes by acting as DNA-binding protein (Ronai et al. 1992). Aldolase is induced under hypoxia that may result from abiotic stress (Kelley and Freeling 1984). Aldolase is linked with tonoplast for the activity of V-ATPase in salt-stressed *Mesembryanthemum crystallinum* that results in sodium ion accumulation in vacuole as a defense strategy (Barkla et al. 2009). Fructose bisphosphate aldolase is speculated in integration of signals linked to the growth, development, and sugar anabolism (Li et al. 2012). In soybean exposed to flooding stress, aldolase protein abundance was increased (Oh and Komatsu 2015). Fructose bisphosphate aldolase is induced by various abiotic stresses in *Arabidopsis* (Lu et al. 2012). The enzyme is also involved in plant development, metabolism and abiotic stress responses (Lv et al. 2017). In the current study, increased protein abundance of aldolase depicts increased rate of glycolysis under flooding stress as plant had limited means to generate energy due to blockage of oxidative phosphorylation.

In the current study, protein abundance of the enolase was increased under flooding stress. The enzyme activity changes also followed the pattern of increase. The enzyme enolase which is also called phosphopyruvate hydratase is an important enzyme of glycolysis, responsible for conversion of 2-phosphoglycerate to phosphoenol pyruvate that ultimately leads to pyruvate formation along-with energy generation. Enolase is induced in maize under anaerobic conditions (Lal et al. 1998). Enolase has also been shown linked to the tonoplast for enabling V-ATPase activity (Barkla et al. 2009). Increase in enolase abundance has been reported in soybean facing flooding stress (Oh and Komatsu 2015; Yasmeen et al. 2016). The results of the present study are in agreement with previous reports indicating that enolase as glycolytic enzyme might have helped in increasing frequency of glycolysis for generating energy under flooding stress.

Alcohol dehydrogenase 1 protein abundance was highly increased under flooding stress as compared to age-matched control plants. Under anaerobic conditions such as flooding, plants ferment glucose to ethanol in the presence of alcohol dehydrogenase. Fermentation thus produces small amount of ATP for life continuity along-with glycolysis (Gibbs and Greenway 2003). Proteomic and transcript abundances of alcohol dehydrogenase are highly increased in soybean under flooding stress (Komatsu et al. 2010b; Komatsu et al. 2011; Oh and Komatsu 2015). Activities of alcohol dehydrogenase were remarkably increased in soybean leaf under flooding stress (Wang et al. 2017). From the previous reports as well as results of current study, the evidence of alcohol dehydrogenase induction and shifting of metabolism to anaerobic mode is confirmed. Soybean used anaerobic fermentation to increase its ATP for survival under flooding stress.

Plant invertase/pectin methylesterase inhibitor was increased in protein abundance and activity. The enzyme activity was much higher when measured at the end of 3^rd^ and 4^th^ day of flooding stress. Pectin plays roles in controlling cell wall porosity (Braybrook et al. 2012), cell adhesion Dahir and Braybrook 2015) and a key factor in plant development (Levesque-Trembley et al. 2015; Saffer 2018). Pectin methylesterase (PME) brings esterification. The extent of methylesterification determines the susceptibility of the plant cell wall to the pectin-degrading enzymes (Lionetti et al. 2012). Plant PME activity generates methanol as a signal of the damaged self, leading to regulate the transcription of pathogen-related PME inhibitor (PMEI) genes (Lionetti et al. 2017). Studies suggest that inhibitory activities of PMEIs are crucial depending on the cell wall environment and different specificities for target PMEs for ensuring a development- and/or stress-dependent adjustments in cell wall (Wormit and Usadel, 2018). Plant invertase/PMEI abundance and/or activity increased in soybean under flooding stress in current study as well as previous findings by Oh and Komatsu (2015) and Yasmeen et al. (2016). These reports suggest that cell wall brings re-adjustments in its structure and mechanics as a mechanism to deal with the flooding stress.

## Conclusions

Flooding acts as abiotic stress for soybean that brings hypoxic or anoxic conditions on the plant. Soybeans respond to flooding stress by altering its basic metabolic modes. It restricts the normal metabolism and brings reduction in ATP yielding and high energy consuming processes. Plant accelerates glycolysis as glycolytic enzymes such such as aldolase, enolase etc. increase their protein abundances and activities. Side-wise, after glycolysis, pyruvate undergoes fermentation pathway to yield ethyl alcohol. Multi-faceted Cupins and toxics scavenging glyoxalases also play cruicial roles in stress responses. Cell wall being outer boundary of plant cell is at high exposure to flooding stress but brings alterations and rearrangements in its structure and mechanics through vaious enzymes such as pectin methylesterase inhibitors to cope with the flooding stress. Thus, soybean brings biochemical and structural changes to effectively respond to flooding stress and adopts the less energy consuming strategies for the survival.

### Supporting information

S1 Table. Sieve MS data of untreated control soybeans. (Excel)

S2 Table. Sieve MS data of flooded soybeans. (Excel)

S3 Table. Sieve MS data of 364 commonly identified proteins in control and flooded soybeans. (Excel)

## Acknowledgements

The authors would like to thank the Deanship of Scientific Research at Taif University for funding this work through Taif University Researchers Supporting Project number (TURSP – 2020/141), Taif University, Taif, Saudi Arabia.

## Authors Contributions

M.N.K and I.A designed, performed the experiment and wrote the manuscript draft. I.D, M.T, M.K, E.A and M.I.K edited the manuscript. A.N and H.D provided funds for the research and critically reviewed the manuscript.

